# Neurocognitive and Functional Heterogeneity in Depressed Youth

**DOI:** 10.1101/778878

**Authors:** Erica B. Baller, Antonia N. Kaczkurkin, Aristeidis Sotiras, Azeez Adebimpe, Danielle S. Bassett, Monica E. Calkins, Zaizu Cui, Raquel E. Gur, Ruben C. Gur, Kristin A. Linn, Tyler Moore, David. R. Roalf, Erdem Varol, Daniel H. Wolf, Cedric H. Xia, Christos Davatzikos, Theodore D. Satterthwaite

## Abstract

**OBJECTIVE:** Depression is a common psychiatric illness that often begins in youth, and is associated with cognitive symptoms. However, there is significant variability in the cognitive burden, likely reflecting biological heterogeneity. This study sought to identify neurocognitive subtypes in a large sample of depressed youth, and evaluated the neural signatures of these subtypes.

**METHODS:** Participants were drawn from the Philadelphia Neurodevelopmental Cohort, including 712 youth with a lifetime history of a major depressive episode and 712 typically developing (TD) youth matched on age and sex. A subset (n=368, TD=200) also completed neuroimaging. Cognition was assessed with the Penn Computerized Neurocognitive Battery. A semi-supervised machine-learning algorithm, HYDRA (Heterogeneity through Discriminative Analysis), was used to delineate neurocognitive subtypes. Subtypes were evaluated for differences in both clinical psychopathology and brain activation during an *n*-back working memory fMRI task.

**RESULTS:** HYDRA identified three neurocognitive subtypes in the depressed group. Overall, Subtype 1 had better performance than TD comparators across many cognitive tasks (high accuracy, moderate speed), Subtype 2 was cognitively impaired (low accuracy, slow speed), whereas Subtype 3 was impulsive (low accuracy, fast speed). While subtypes did not differ in clinical psychopathology, they diverged in their activation profiles in regions critical for executive function, which mirrored differences in cognition.

**CONCLUSIONS:** Using a data-driven approach, three neurocognitive subtypes of depression were identified that differed in neural signatures despite similar clinical psychopathology. These data suggest disparate mechanisms of cognitive vulnerability and resilience in depression, which may inform the identification of biomarkers for prognosis and treatment response.

## INTRODUCTION

Depressive disorders are common, and are consistently among the leading causes of disability world-wide (1). Major depressive disorder (MDD) is frequently difficult to treat, with one-third of patients remaining symptomatic despite treatment (2). Depression often starts in adolescence, with a 4-5% prevalence of MDD in youth (3). Like adults, there is significant heterogeneity in response to treatment in youth with MDD. The high variability in treatment response suggests that MDD is a heterogeneous illness, with multiple pathophysiologic pathways that converge on a similar clinical phenotype (4). Notably, the *Diagnostic and Statistical Manual of Mental Health Disorders* (DSM-5) relies solely on clinical symptom classification (5).

Though mood symptoms are most assessed in MDD, deficits in cognitive processing are strong predictors of poor outcome. Problems in the domains of planning, executive function, short- and long-term memory, and social cognition have been linked to poor educational attainment, lack of employment prospects, and downward social mobility (6,7). However, similar to treatment-response, there is substantial heterogeneity in cognitive function in depression. Some youths experience profound mood symptoms with cognitive resilience, whereas others experience marked cognitive impairment.

Given their involvement in higher level cognitive processing, the frontoparietal and cingulo-opercular networks have emerged as targets in the study of youth depression. The few studies that have evaluated cognitive control in depressed youth have yielded mixed results. Whereas some studies have shown robust differences in activation in the prefrontal cortex in depressed youth as compared to healthy controls, others have found no significant differences (8,9). In contrast, other studies have shown abnormal anterior cingulate cortex activation in depressed youth (10,11). Of note, none of these studies characterized or evaluated cognitive heterogeneity, which may account for the conflicting findings. Given the high degree of cognitive heterogeneity in depression and the important relationship between cognitive function and functional outcome, we sought to identify neurocognitive subtypes in youths with a history of depression, and uncover neural patterns associated with these subtypes using a data set independent of cognitive clustering.

Machine learning tools are increasingly capable of uncovering homogeneous subtypes within heterogeneous conditions like MDD (12). We used a recently developed semi-supervised machine learning algorithm HYDRA (Heterogeneity through Discriminative Analysis) to identify cognitive subtypes within a sample of youth with a history of a major depressive episode compared to a group of typically developing youth. We then evaluated the cognitively-defined subgroups on independent measures of clinical symptoms and brain activation during an *n*-back working memory task. We predicted that we would identify cognitive subtypes that had distinct neural signatures. In line with previous work, we anticipated that better cognitive performance would be associated with greater brain activation in the *n*-back (13). Given the data-driven approach of this study, we expected that new relationships between cognitive deficits, clinical symptoms, and brain activation would emerge.

## METHODS

### Participants

A total of 9,498 participants aged 8-22 years received cognitive assessment and clinical phenotyping as part of the Philadelphia Neurodevelopmental Cohort (PNC), a large community-based sample of youths (14,15). A randomly selected subsample (n = 1,601) also received multimodal magnetic resonance imaging (MRI). We excluded participants with medical disorders that could impact brain function or missing data (see Supplement for details). Of the remaining participants, 712 met screening criteria for a lifetime history of a major depressive episode as defined by DSM-IV-TR, and 2,310 were typically developing (TD) youth with no psychiatric diagnosis (16). For the sake of clarity, we will refer to youths with a history of a major depressive episode as *depressed youths* (DY). Using the R package MatchIt, depressed youths were matched to individuals in the TD group on the basis of age and sex as detailed in the Supplement (17). Because not all participants recieved neuroimaging, we preferentially included those who completed neuroimaging. This matching procedure yielded a final sample of 712 DYs and and 712 TDs (**Table 1A**). A subset of these youth (DY = 168, TD = 200; **Table 1B**) also completed the *n-*back working memory task during functional magnetic resonance imaging (fMRI) and passed strict quality control criteria as detailed in the Supplement. The institutional review boards of both the University of Pennsylvania and the Children’s Hospital of Philadelphia approved all study procedures.

**Table 1:** Sample demographics for the whole group (A) as well as for the imaging subsample (B). Typically developing and depressed youth were matched on age and sex prior to subtyping.

| Demographics of Whole Group |  |  |  |
| --- | --- | --- | --- |
|  | Typically Developing | Depressed | *p* |
| n | 712 | 712 |  |
| % Caucasian | 64 | 55 | 0.001 |
| % Female | 67 | 67 | 1.00 |
| Maternal Education (mean[sd]) | 14.93 [2.52] | 14.12 [2.29] | <0.001 |
| Age (mean[sd]) | 16.21[2.91] | 16.13 [2.90] | 0.94 |

**B**
| Demographics of Imaging Subsample |  |  |  |
| --- | --- | --- | --- |
|  | Typically Developing | Depressed | *p* |
| n | 200 | 168 |  |
| % Caucasian | 61 | 46.4 | <0.001 |
| % Female | 59 | 64 | 0.34 |
| Maternal Education (mean[sd]) | 14.93 [2.58] | 13.98 [2.36] | <0.001 |
| Age (mean[sd]) | 16.49[2.84] | 16.87 [2.20] | 0.57 |

### Clinical assessment and clinical factor analysis

As described in detail in the Supplement, assessment of lifetime psychopathology was conducted using GOASSESS, a structured screening interview based on a modified version of the K-SADS (18). We included those who met screening criteria for a lifetime history of a major depressive episode, which included both symptoms of depression and resulting distress and impairment. To provide a dimensional summary of the diverse psychopathology data for all 1,424 youths, we used a confirmatory bifactor analysis to orthogonally model four factors (anxious-misery, psychosis, externalizing, and fear) plus a general factor, overall psychopathology, which represents the symptoms common across all psychiatric classifications (19). To avoid analytic circularity, our factor analysis excluded all items from the depression section of the interview that were used to identify participants with a history of depression.

Given that anxiety is frequently comorbid with depression and has many overlapping symptoms, we evaluated measures of anxiety separately. Since the depression group was classified based on a lifetime history of depression irrespective of current mood state, but mood state may impact cognition, we also evaluated state and trait anxiety scores at the time of MRI using the State-Trait Anxiety Inventory (STAI). Previous work with the STAI has established that it taps a dimensional measure of anxious-misery (20,21).

### Cognitive assessment

Cognition was assessed using the University of Pennsylvania Computerized Neurocognitive Battery (CNB) (22). Twenty-six measures obtained from 14 neurocognitive tests of performance were assessed (14 for accuracy, 12 for speed). Domains included executive functioning (3 tests), episodic memory (3 tests), social cognition (3 tests), complex reasoning (3 tests), and sensorimotor speed (2 tests) as detailed in and the Supplement. Verbal intelligence was estimated with the *Wide Range Achievement Test, 4*^*th*^ *Edition* (WRAT-4) reading subscale with total subscale scores reported as T-scores (mean=100, SD=15) (23).

### Parsing cognitive heterogeneity with semi-supervised machine learning

To identify cognitive subtypes within depressed youths, we used a recently developed semi-supervised machine learning tool: HYDRA (Heterogeneity through Discriminative Analysis). HYDRA compares a reference group (e.g. controls) to a target group (e.g. patients) to identify *k* subtypes (clusters) within the target group (24). In contrast to fully-supervised learning techniques, which cannot distinguish between subtypes of patients, HYDRA simultaneously performs classification and clustering (**Figure 1A**). In contrast to unsupervised clustering techniques (such as *k*-means or community detection), this semi-supervised algorithm clusters the differences between the two groups, rather than clustering the groups themselves, thereby parsing phenotypic heterogeneity of underlying neurobiological processes. To accomplish this, HYDRA finds multiple hyperplanes, which together form a convex polytope that separates controls from subtypes. Rather than coercing participant data points into a single common discriminative pattern, HYDRA allows for the separation of distinct groups distinguished by multiple decision boundaries. The result is a data-driven approach to identifying subtypes of depressed youth that can be further evaluated on independently-measured clinical and imaging characteristics.

**Figure 1:**
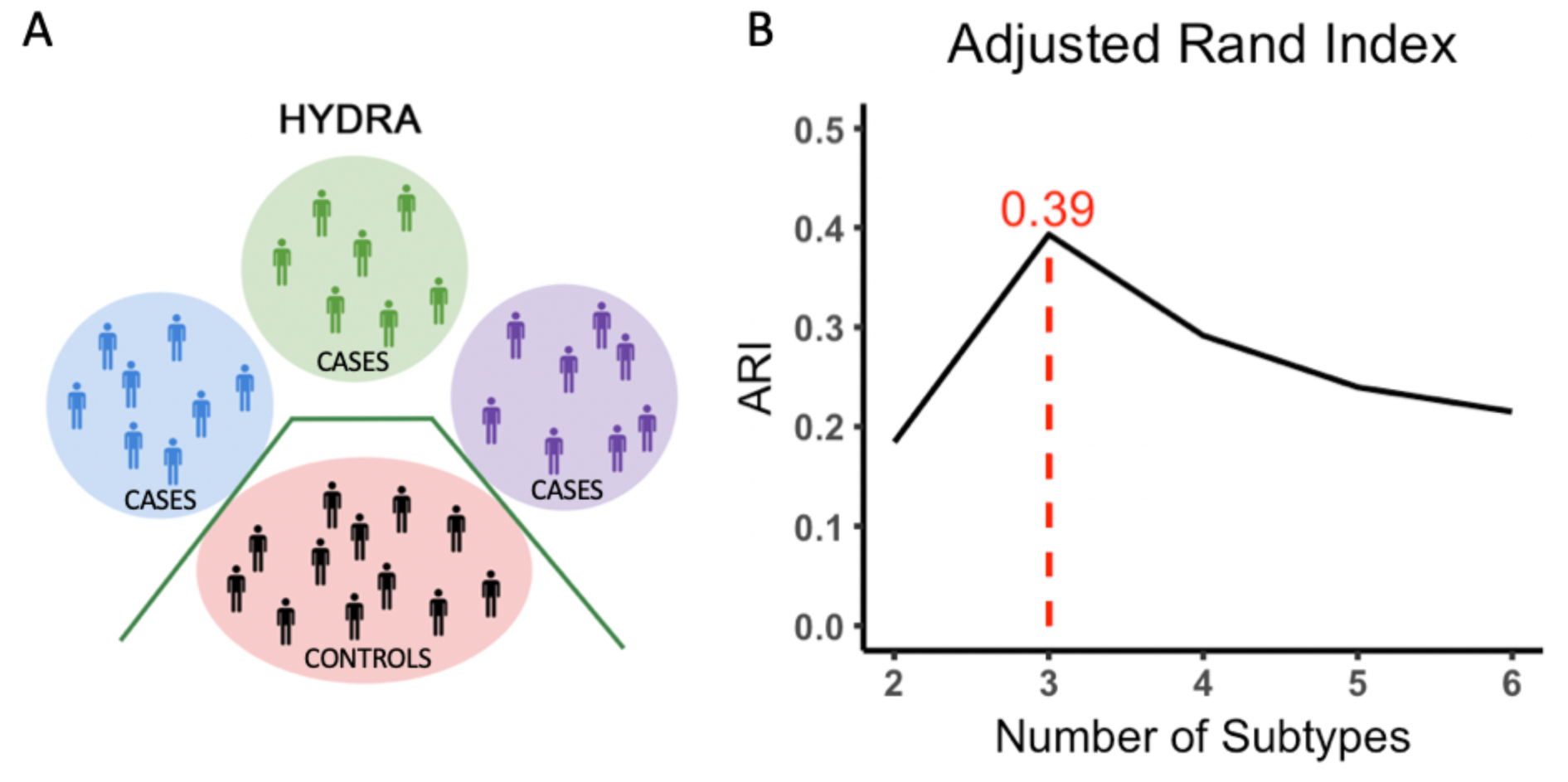
Heterogeneity through Discriminative Analysis (HYDRA) algorithm and subtype selection. (**A**) HYDRA is a semi-supervised machine learning algorithm that reveals homogenous subtypes within a clinical group by maximizing subtype-specific margins between patient clusters and controls, while adjusting for covariates and determining cluster memberships, as shown schematically here. (**B**) The stability of the clustering solution after cross-validation was evaluated over a resolution range of 2-10 clusters (2-6 shown here), and was quantified by the adjusted rand index (ARI). The best clustering solution with the highest ARI is with three subtypes.

HYDRA was used to define cognitive subtypes using the 26 cognitive measures of accuracy and speed. Given known developmental and sex differences in cognition, both age and sex were included as covariates in HYDRA. Consistent with prior studies using this technique, we derived multiple clustering solutions requesting 2 to 10 clusters in order to obtain a range of possible solutions (24,25). The adjusted Rand index (ARI) was calculated using 10-fold cross validation to evaluate the stability of each solution; the solution with the highest ARI value was selected for subsequent analyses.

### Image acquisition, quality assurance, and image processing

Task paradigm, image acquisition, and preprocessing methods are as previously detailed (26) and described in the Supplement. A single scanner (Siemens 3-T Tim Trio) was used to acquire structural and functional MRI data for all subjects. A fractal version of the *n*-back task was used to probe working memory function. Time-series analysis of subject-level imaging data used the FMRIB Software Library version 5.0 (FSL) to model three condition blocks (0-back, 1-back, and 2-back). Subject-level statistical maps were corrected for distortions, co-registered to the structural image using boundary-based registration, and normalized to a custom 1mm adolescent template using Advanced Normalization Tools (27). All transformed images were concatenated so that only one interpolation was performed, and down-sampled to 2mm^3^ voxels prior to analysis. The mean percent signal change on the primary contrast of interest (2-back vs. 0-back) was extracted from 21 previously-defined regions of interest (ROIs) within the executive system (**Supplement Figure 1**) (13).

### Group-level statistical analyses

Having identified subtypes of depressed youth, we sought to understand the characteristics of these subtypes. As our subtypes were defined using cognitive performance data, we first sought to assess the cognitive profiles of each subtype. Notably, statistical testing of cognitive performance between subtypes was not performed, because the cognitive data was used in the clustering procedure and therefore subtypes differed in cognitive performance by definition.

In contrast, clinical symptomatology and neuroimaging were independent data types that were not used in the clustering procedure, and thus were amenable to statistical testing. Accordingly, as a first step we evaluated the clinical profiles of subtypes and controls. Finally, we evaluated whether subtypes displayed differential brain activity in the *n*-back working memory task within the 21 executive system ROIs.

For all analyses, we used a general linear model to test how well the subtype identification predicted the outcome of interest (clinical or imaging measures). In the model evaluating differences in activation during the *n*-back task, we included mean in-scanner motion as an additional model covariate to control for the potentially confounding effects of motion on image quality. Our group variable was modeled as a factor variable, including the typically developing group and the cognitive subtypes. An omnibus ANOVA testing for group differences was corrected for multiple comparisons by controlling the False Discovery Rate (FDR, *Q* < 0.05). For measures that passed FDR correction, we then conducted pairwise *post hoc* tests to determine which subtypes significantly differed from each other; these *post hoc* tests were corrected for multiple comparisons using the Tukey method. Age-by-sex interactions in the ROIs were evaluated separately and all between group differences were found to be not significant. All statistical analysis were performed in R (version 1.1.383).

### Data and code availability

See https://github.com/PennBBL/baller_heterogen_2019 for an overview and all data analysis code used in this manuscript. Data from the Philadelphia Neurodevelopmental Cohort can be accessed at https://www.ncbi.nlm.nih.gov/projects/gap/cgi-bin/study.cgi?study_id=phs000607.v3.p2. The HYDRA code can be found at https://github.com/evarol/HYDRA.

## RESULTS

Of the ten possible clustering solutions generated by HYDRA, a well-defined peak at *k* = 3 (ARI = 0.39) emerged, suggesting the presence of three distinct neurocognitive subtypes of depressed youth (**Figure 1B**). As an initial step, we evaluated the demographics of the neurocognitive subtypes, and found that the subtypes did not differ in age or sex. However, Subtype 2 had a lower percentage of Caucasians and lower maternal education level. While significant, this difference was relatively modest: on average, mothers had some college education, and differed at most by approximately 1.5 years (Subtype 1 vs. Subtype 2).

We next characterized subtypes on their overall patterns of accuracy and speed (**Figure 2A**). Across all accuracy domains, Subtype 1 consistently outperformed both other depressed subtypes as well as the TDs (**Figure 2B**). In contrast, when cognitive speed was evaluated, Subtype 1 performed similarly to the TDs, with faster speed than Subtype 2 and slower speed than Subtype 3 (**Figure 2C**). Overall, Subtype 1 was a high-performing subset of depressed participants, who were able to efficiently maximize the trade-off between accuracy and speed. In contrast to the high-performing Subtype 1, Subtype 2 showed globally impaired cognition, with the lowest accuracy and slowest speed of all subtypes. Finally, Subtype 3 had poor accuracy performance but very fast speed, suggesting that Subtype 3 was impulsive, and was unable to accurately balance the competing demands of accuracy and speed.

**Figure 2:**
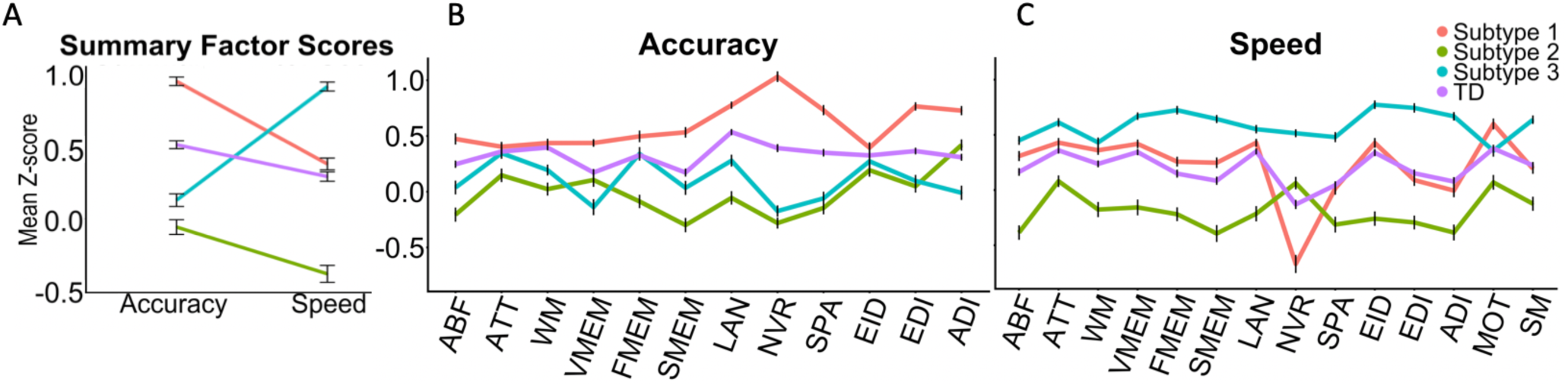
Subtypes revealed by HYDRA differ in neurocognitive profile. (**A**) Three neurocognitive signatures emerged in depressed youth: Cluster 1 had preserved cognition, with high accuracy and speed; Cluster 2 had impaired cognition, with low accuracy and speed; Cluster 3 was impulsive, with high speed but low accuracy. (**B-C**) Patterns were largely consistent for all measures of accuracy (panel **B**) and speed (panel **C**) (*P*_*fdr*_ <0.05 for all measures). HYDRA-Heterogeneity through Discriminative Analysis, ABF-Abstraction/Mental Flexibility, ATT-Attention, WM-Working Memory, VMEM-Verbal Memory, FMEM-Face Memory, SMEM-Spatial Memory, LAN-Language/Verbal Reasoning, NVR-Nonverbal Reasoning, SPA-Spatial Reasoning, EID-Emotion Recognition, EDI-Emotion Discrimination, ADI-Age Discrimination, MOT-Motor, SM-Sensorimotor

We next evaluated the differences in clinical symptom profiles across TDs and all subtypes by evaluating dimensions of psychopathology that were not used during clustering. As expected, all subtypes had increased psychopathology compared to TDs across most dimensions, including anxious-misery, externalizing, fear, and overall psychopathology (*P*_*fdr*_ < 0.05). The psychosis factor did not differ across TDs and subtypes. Despite such clear differences from controls, there were very few significant differences in clinical symptoms between the subtypes (**Supplement Table 1**). Across the clinical measures evaluated, the subtypes only differed on the fear dimension (Subtype 2 > Subtype 1, *p <* 0.0001; Subtype 2 > Subtype 3, *p* <0.0001). Subtype 1 also had slightly more anxious-misery than Subtype 2 (*p*=0.03). Similarly, there were no differences between the neurocognitive subtypes in state or trait measures of anxiety (**Supplement Table 2)**, indicating that the neurocognitive subtypes did not simply reflect the underlying clinical symptomatology.

As a final step, to test the hypothesis that neurocognitive subtypes reflected distinct neural profiles, we examined the between-group differences in *n-*back working memory activation. We specifically compared mean percent signal change in 21 *a priori* ROIs within the executive system (13). Of these 21 regions, six showed significant group differences (*P*_*fdr*_ < 0.05; **Figure 3A**): left anterior dorsolateral prefrontal cortex, anterior cingulate, left dorsal frontal cortex, bilateral precuneus, and right crus II. Differences in activation mirrored the neurocognitive profiles of the subtypes. Specifically, high-performing Subtype 1 had globally higher activation, even exceeding that of TDs. Similarly, Subtype 2 had the lowest activation, while Subtype 3 showed reduced activation that was less severely impaired than Subtype 2 (**Supplement Table 3**). In summary, neurocognitive subtypes had distinct neural signatures that scaled with cognitive performance, despite the similar clinical symptomatology of these subtypes.

**Figure 3:**
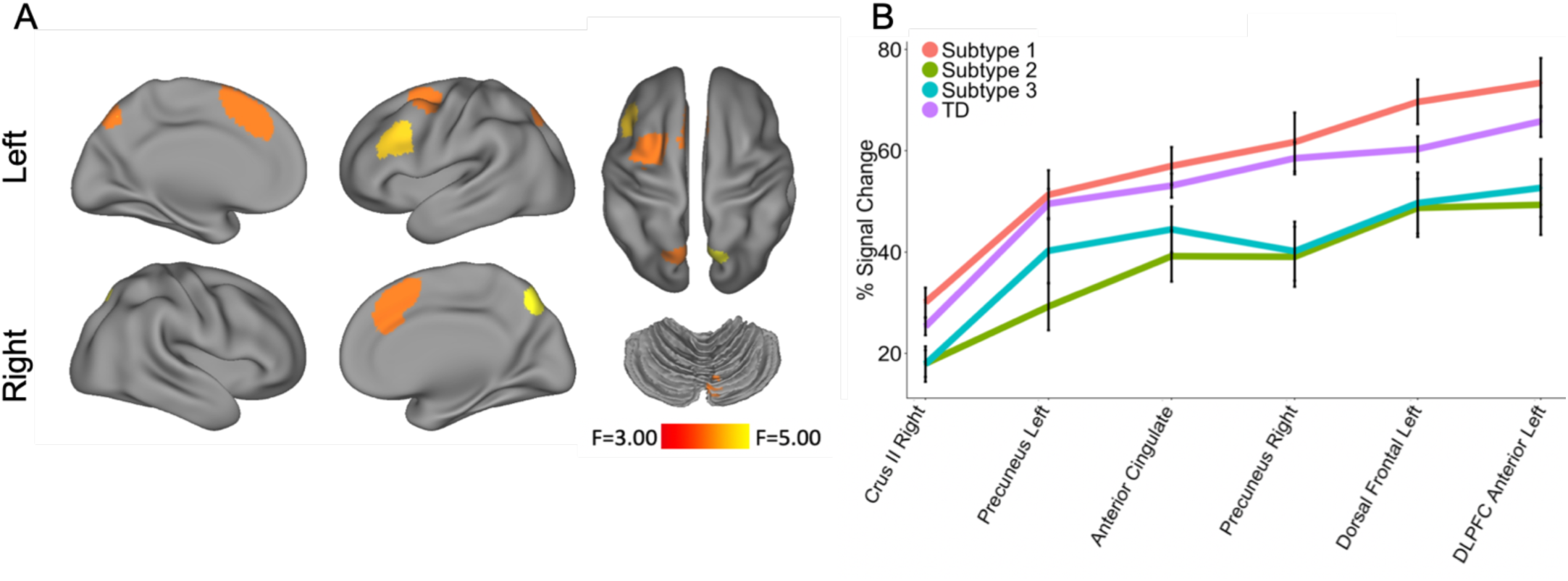
Neurocognitive subtypes differ in activation of executive regions during an *n*-back working memory paradigm. (**A**) Group differences (*P*_*FDR*_ <0.05) in *n*-back activation between subtypes were present in six functionally defined regions of interest previously described (13). See Supplementary Figure 1 for all twenty-one pre-defined regions of interest. (**B**) Group differences were driven by a consistent pattern across regions, with greater activation in Subtype 1 and TDs than in Subtype 2 or Subtype 3.

## DISCUSSION

Using a recently-developed semi-supervised machine learning algorithm and a large sample of youth with a history of depression, we identified three distinct neurocognitive subtypes of depression. Subtype 1 had globally preserved cognition, and outperformed the typically developing youth on all domains. Subtype 2 had globally impaired cognition, while Subtype 3 was impulsive, sacrificing accuracy for speed. The activation profiles of each subtype during the *n-*back task paralleled the neurocognitive signatures. This concordance between cognitive and neuroimaging results implies that our data-driven approach identified biologically-relevant subtypes. Importantly, these subtypes were not clearly distinguishable based on their clinical symptoms. The discrepancy between subjective clinical symptoms and objective cognitive and neural measures is intriguing given that psychiatric illnesses and treatment recommendations are currently based solely on observed clinical symptoms. Our study highlights both the important heterogeneity of cognitive dysfunction in depression, and the promise of machine learning for understanding this and other types of heterogeneity in psychiatric disorders.

All functionally-defined regions of interest evaluated in this study are consistently recruited by working memory load and play a central role in executive functioning (26). Notably, the subset of regions that showed between-subtype differences map onto the frontoparietal and cingulo-opercular networks, which are of particular developmental relevance. The frontoparietal network balances cognitive flexibility with cognitive control, both within the network as well as by coordinating network activity in separate distributed networks (28,29). Throughout healthy adolescent brain development, the frontoparietal network increases in strength and flexibility (30). Dysfunctional development of this network, on the other hand, appears to be a risk factor for psychopathology (26,31). Brain imaging studies in people with affective disorders, many of whom have their onset in adolescence, tend to show abnormalities in activity in the frontoparietal network (32). In our study, youths with preserved cognition had consistently higher activation in the frontoparietal regions, with both higher cognitive performance and greater activation as compared to typically developing youth. Both the low-functioning and impulsive groups had lower activity in the frontoparietal regions, suggesting that both poor overall performance and inability to exert control of the speed/accuracy trade-off were due to under-functioning of this network.

The cingulo-opercular network, which regulates salience and inhibitory control, is similarly important in development and in affective disorders, and also showed differences between the subtypes. Typically developing adolescent brains show progressive strengthening of the cingulo-opercular network, reflecting the ability to process salient information and to engage in impulse control when selecting behaviors (33,34). Abnormal functioning of this system has been associated with anhedonia in youth as well as attention-deficit hyperactivity disorder (31,35). Our findings similarly showed neurocognitive subtype-specific differences in a hub of the cingulo-opercular network, the dorsal anterior cingulate, which is less active in youths with poor overall cognition or high impulsivity.

Research in adults suggests that both cognition and neural activity can serve as markers for prognosis and treatment in depression. Higher baseline cognitive functioning, specifically in the domains of executive function and working memory, predicts better response to selective serotonin reuptake inhibitors (36). The left dorsolateral prefrontal cortex is of particular relevance because it is the most common target of TMS therapy (37,38)9/22/19 10:54:00 AM. Studies of the neural correlates of cognitive behavioral therapy reported that increased activity in the left and right precuneus (part of the frontoparietal network) predict symptom improvement (39). Neural activity in these brain regions also significantly differed by subtype in our study. Our findings suggest that neurocognitive heterogeneity is prominent in depressed youth and is mirrored by underlying neural heterogeneity. As in prior studies of adults, targeting the cognitive system or the underlying neural networks with behavioral, pharmacological, or neuromodulatory treatment may likewise prove useful for the treatment of youth with depression.

Despite clear differences in cognition and neural activity in the neurocognitive subtypes, these differences did not map clearly onto clinical symptom differences. The lifetime clinical symptom profiles as well as anxious-misery symptoms (assessed by the STAI) on the date of testing were largely similar between subtypes, indicating that the cognitive and neural differences observed between subtypes did not merely reflect differences in clinical presentation. This pattern of results aligns with data suggesting that patients with a similar symptomatic presentation may have divergent cognitive profiles, prognosis, and response to treatment (36). Furthermore, this result stands in contrast with previous studies that have identified subtypes of depression based on the presence or absence of anxious-misery (12,40).

This study adds new insights to the growing body of research that uses machine learning to understand heterogeneity in psychiatry. Previous studies have primarily used either unsupervised machine learning algorithms or supervised machine learning algorithms, both of which have inherent limitations (12,35). Unsupervised machine learning algorithms allow subjects to be clustered into subtypes, but do not account for important data like clinical diagnosis. In contrast, supervised machine learning algorithms can be used to classify a control group from a patient group based on imaging or other data, but cannot assess heterogeneity. Our study leverages both techniques by using a semi-supervised machine learning algorithm that simultaneously performs classification (i.e. control vs. case) and clustering (subtypes of cases). This approach resembles real-world decision-making, whereby a practitioner must be able to distinguish a depressed person from a healthy person, and also simultaneously attempt to stratify the individual into meaningful subtypes that inform management.

Several limitations should be noted. First, due to the cross-sectional nature of the data, we cannot link our data-driven subtypes to meaningful functional outcomes. Since some youth with depression go on to have long-term functional limitations in a variety of domains, it would be important to consider cognitive vulnerability early. Our study was also limited by an assessment that evaluated only a lifetime history of a major depressive episode, but did not evaluate whether participants were currently depressed. However, our state measures of anxious-misery were not different between subtypes, suggesting that there is a low likelihood that current affective symptoms drove the observed differences between subtypes.

In sum, capitalizing upon a large sample of youth with a history of depression and recently-developed machine learning tools, we delineate novel neurocognitive subtypes of depression in youth. These results have the potential to inform longitudinal studies that could describe how cognitive heterogeneity in depression impacts disease progression and functional outcomes. Likewise, understanding how heterogeneous cognitive and neural deficits moderate treatment response is a critical next step. Finally, next-generation personalized medicine approaches could deploy neuromodulatory therapies that may be particularly useful for subtypes of depressed youth with impaired cognitive function and distinct neural deficits.

## ACKNOWLEDGEMENTS

This work was supported by grants from the National Institute of Mental Health (NIMH; grant numbers: R01MH120482, R01MH107703, R01MH112847 and R01MH113550 to TDS; K99MH117274 to ANK; R01MH107235 to RCG; R01MH13565 to DHW; and R01MH11207 to CD. Additional support was provided by the Lifespan Brain Institute at the Children’s Hospital of Philadelphia and Penn Medicine. The PNC was funded by RC2 grants MH089983 and MH089924 to REG from the NIMH. Support for developing multivariate pattern analysis software (AS & TDS) was provided by a seed grant by the Center for Biomedical Computing and Image Analysis (CBICA) at Penn. Support was also provided by a NARSAD Young Investigator Award (ANK) as well as a Penn PROMOTES Research on Sex and Gender in Health grant (ANK) awarded as part of the Building Interdisciplinary Research Careers in Women’s Health (BIRCWH) grant (K12 HD085848) at the University of Pennsylvania.

