## Supplementary Materials for "Neurocognitive and Functional Heterogeneity in Depressed Youth"

**Table of Contents**


**Participants** 2

**Matching Procedure** 2

**Clinical Assessment** 3

**Cognitive Assessment** 4

**Clinical Factor Analyses** 4

***n*-back task** 5

**Image Acquisition and preprocessing** 5

- **Acquisition** 5
- **Preprocessing** 6
- **Construction of functional regions of interest** 7

**References** 8

**Supplementary Tables and Figures** 9

- **Supplementary Table 1: Post hoc analyses of clinical factor scores** 9
- **Supplementary Table 2: Post hoc analyses of state and trait anxiety** 10
- **Supplementary Table 3: Post hoc analyses of *n*-back activation** 11
- **Supplementary Figure 1: Twenty-one functionally defined regions of interest.** 12

**Participants:**

A total of 9,498 participants 8-22 years of age received cognitive assessment and clinical phenotyping as part of the Philadelphia Neurodevelopmental Cohort (PNC), a large community-based sample of youth (1). From the pool of 9,498 participants, 6,476 were ineligible for the current study for 1) medical disorders that could impact brain functioning (n = 2,347), 2) missing age at clinical assessment (n = 92), or 3) missing depression or overall psychiatric sub-score (n = 87); several subjects were ineligible based on multiple criteria. Of the remaining participants, 712 met screening criteria for a lifetime history of a major depressive episode (referred to as depressed youth, or DY) and 2,310 were typically developing (TD) youth with no psychiatric diagnosis. Our analysis evaluated a final sample of 712 depressed youth and 712 typically developing youth (total n = 1,424).

Of the 1,424 individuals in the final group, a subset of participants (n=368, TD=200) also completed *n*-back functional magnetic resonance imaging (fMRI) and passed strict quality control including T1 structural and motion exclusion.

**Matching Procedure:**

Using the R package Matchit, depressed youth were age and sex matched with typically developing youth. Given that not all participants underwent neuroimaging, the match was performed in multiple steps to allow us to enrich our typically developing group for children who were in the subset that obtained neuroimaging. All depressed youth were included in the final sample. First, depressed youth with imaging were matched with TD youths with imaging. Next, depressed youth without imaging were matched with the remaining TDs with imaging that were not matched in the first step. Youths with poor T1 quality were excluded. The results from both matches were combined, yielding a group of 712 DYs and 712 TDs. After additional quality assessment of the *n*-back task-related imaging data, 368 youth (DY =168 , TD = 200) were included in the functional imaging analysis.

**Clinical Assessment**:

As described in detail in our previous work, assessment of lifetime psychopathology was conducted using GOASSESS, a structured screening interview administered to probands (age 11-22 years) and collateral informants of probands (age 8-17 years), based on a modified version of the Kiddie-Schedule for Affective Disorders and Schizophrenia and Diagnostic and Statistical Manual of Mental Disorders, 4th edition, Text Revision criteria (2). The GOASSESS interview assesses lifetime occurrence of mood (major depressive episode, mania), anxiety (agoraphobia, generalized anxiety, panic, specific phobia, social phobia, separation anxiety, posttraumatic stress), behavioral problems (oppositional defiant, attention deficit/hyperactivity, conduct), psychosis, eating disorder (anorexia, bulimia), and suicidal symptoms. Among the GOASSESS questions, 107 screening items administered to all participants were used for the current investigation. Of note, due to comorbidity, participants may be represented in more than one diagnostic category. The GOASSESS interview was administered by trained assessors who underwent a common training protocol (developed and implemented by MEC) that included didactic sessions, assigned readings, and supervised pair-wise practice. Assessors were certified for independent assessments following observation by a certified clinical observer who rated the proficiency of the assessor on a 60-item checklist of interview procedures. The median interval of time between clinical assessment and neuroimaging was 2 months.

**Cognitive Assessment:**

Cognition was assessed using the University of Pennsylvania Computerized Neurocognitive Battery (CNB), which has been described previously (3). Briefly, 14 cognitive tests evaluating aspects of cognition, including executive control, episodic memory, complex reasoning, social cognition, and sensorimotor speed, were administered in a fixed order. Except for two sensorimotor tests that only measure speed, each test provides measures of both accuracy and speed, yielding 26 total measures (abstraction/mental flexibility, attention, working memory, verbal memory, face memory, spatial memory, language/verbal reasoning, nonverbal reasoning, spatial reasoning, emotion recognition, emotion discrimination, age discrimination, motor, sensorimotor). Academic skills were measured with the Wide Range Achievement Test, 4th Edition (WRAT-4) reading subscale with total subscale scores reported as T-scores. Youth performance on each measure was transformed into a Z-score, which was used for further analysis.

**Clinical Factor Analyses:**

To provide a dimensional summary of the diverse psychopathology data, we applied an exploratory factor analysis to 107 item-level symptom questions from the GOASSESS interview, which has been described in detail elsewhere (3). Of note, contrary to the previous published analysis which utilized 112 item-level symptoms questions, we excluded 5 depression items that were used to define our diagnostic groups. This exploratory factor analysis yielded four correlated dimensions of psychopathology including factors for anxious-misery (31 items), psychosis (26 items), behavioral (externalizing, 25 items), and fear (25 items). We then used a confirmatory bifactor analysis implemented in Mplus11 to orthogonally model the four factors plus overall psychopathology, which represents the symptoms common across all psychiatric disorders. Given that some anxiety sub-scores were used in constructing the depression summary score, the factor analysis presented in this paper excluded measures that were used for the diagnosis of a major depressive episode.

***n*-back task:**

Subjects completed a fractal version of the *n*-back task during their fMRI scan (4,5). During the task, a fractal was presented for 500 ms followed by a 2500 ms interstimulus interval. This task was used to probe working memory and had 3 conditions: 0-, 1-, and 2-back. During the 0-back, subjects responded by pressing a button when the fractal presented matched a predefined fractal. During the 1-back condition, subjects responded when the fractal presented was the same as the one preceding it. During the 2-back condition, subjects responded when the fractal was identical to the one two before it. Each condition consisted of three 20-trial blocks, each preceded by a 9s instruction period, with a target to foil ratio of 1:3. The task included a total of 45 targets and 135 foils, as well as three 24 s blocks of rest during which a fixation crosshair was displayed.

**Image Acquisition and preprocessing:**

*Acquisition*

Imaging data were acquired on a single 3T Siemens TIM Trio whole-body scanner using a 32-channel head coil. A magnetization-prepared rapid acquisition gradient echo T1-weighted (MPRAGE) image (TR, 1810 ms; TE, 3.51 ms; TI, 1100 ms; FOV, 180 × 240 mm; matrix, 192 × 256; 160 slices; slice thickness/gap, 1/0 mm; flip angle, 9°; effective voxel resolution, 0.9 × 0.9 × 1 mm) and B0 field map (TR, 1000 ms; TE1, 2.69 ms; TE2, 5.27 ms; 44 slices; slice thickness/gap, 4/0 mm; FOV, 240 mm; effective voxel resolution, 3.8 × 3.8 × 4 mm) were acquired to aid spatial normalization to standard space and application of distortion correction procedures, respectively. Functional images were then obtained using a whole-brain, single-shot, multislice, gradient-echo echoplanar sequence (231 volumes; TR, 3000; TE, 32 ms; flip angle, 90°; FOV, 192 × 192 mm; matrix 64 × 64; 46 slices; slice thickness/gap 3/0 mm; effective voxel resolution, 3.0 × 3.0 × 3.0 mm).

*Preprocessing*

As previously described, fMRI data were pre-processed with FSL, including skull removal with BET, slice time correction, motion-correction with MCFLIRT, spatial smoothing (6 mm FWHM), and mean-based intensity normalization. Subject-level timeseries analyses were carried out using FILM3 (FMRIB's Improved Linear Model) with local autocorrelation correction (5). The three condition blocks (0-back, 1-back, and 2-back) were modeled using a canonical (double-gamma) hemodynamic response function, with six motion parameters and the instruction period included as nuisance covariates. The rest condition served as the unmodeled baseline. The median functional and anatomical volumes were co-registered using boundary-based registration with integrated distortion correction using FUGUE. The anatomical image was normalized to a custom 1mm template using the top-performing diffeomorphic SyN registration of ANTS (6). All transformed images (distortion correction, co-registration, normalization, and down-sampling to 2mm$^3$) were concatenated so that only one interpolation was required. The statistical maps for the contrast of interest (2-back > 0-back) were then used in the group-level analyses. This contrast was implemented using the task module of XCP (7).

*Construction of functional regions of interest*

Functional regions of interest were delineated from the 2-back > 0-back map from the complete subsample of youth who underwent *n*-back imaging and met quality control (n = 951) as previously described (4). Specifically, to isolate core regions of the executive network with a high degree of anatomic specificity, the 2-back>0-back map was thresholded at z > 20; clusters of <100 voxels were discarded. This high threshold was selected because at lower thresholds substantial volumes of white matter were included due to spatial smoothing, the very high statistical power of the large sample, and the robust nature of the contrast. Next, a watershed algorithm implemented in MATLAB was applied to parse confluent regions of interest. The watershed procedure separates contiguous regions of voxels into subregions by first identifying local maxima, each of which becomes a peak within a subregion. Subregion boundaries are defined by the watershed algorithm, which computes how water would drain into the inverted topology of the activation map. Last, a second threshold on spatial extent (k < 50 voxels) was used to remove undesirably small subregions by absorbing them into the nearest neighboring suprathreshold subregion. When this procedure was applied to the activated contrast of 2-back > 0-back, a set of 21 functional ROIs was produced within the executive network (see Results) that corresponded to a high degree with previously published meta-analyses of working memory (8,9). Finally, signal change in the 2-back > 0-back contrast in each of these 21 regions of interests was extracted. The final regions of interest included the right and left crus I, right and left crus II, right and left dorsolateral prefrontal cortices (anterior region), right and left dorsolateral prefrontal cortices (posterior region), right and left dorsal frontal gyri (part of middle frontal gyrus), right and left precuneus, right and left thalamus, right and left insula, right and left frontal poles, right and left parietal cortices, and dorsal anterior cingulate gyrus (Supplementary Figure 2).

### Supplementary Tables and Figures

**Supplementary Table 1:** *Post hoc* analyses of clinical factor scores. Group statistics were corrected for multiple comparisons by controlling the False Discovery Rate ($Q$<0.05). Pairwise contrasts are reported as $p$-values, and were adjusted via the Tukey method.

| | Pr(>F) | TD vs. Subtype 1 | TD vs. Subtype 2 | TD vs. Subtype 3 | Subtype 1 vs. Subtype 2 | Subtype 1 vs. Subtype 3 | Subtype 2 vs. Subtype 3 |
|---|---|---|---|---|---|---|---|
| Anxious-Misery | <0.001 | <0.001 | <0.001 | <0.001 | 0.030 | 0.375 | 0.724 |
| Externalizing | <0.001 | <0.001 | <0.001 | <0.001 | 0.701 | 0.776 | 0.207 |
| Fear | <0.001 | 0.017 | <0.001 | 0.041 | <0.001 | 1 | <0.001 |
| Overall Psychopathology | <0.001 | <0.001 | <0.001 | <0.001 | 0.313 | 0.957 | 0.671 |

**Supplementary Table 2:** *Post hoc* analyses of state and trait anxiety. Group statistics were corrected for multiple comparisons by controlling the False Discovery Rate ($Q$<0.05). Pairwise contrasts are reported as $p$-values, and were adjusted via the Tukey method.

| | Pr(>F) | TD vs. Subtype 1 | TD vs. Subtype 2 | TD vs. Subtype 3 | Subtype 1 vs. Subtype 2 | Subtype 1 vs. Subtype 3 | Subtype 2 vs. Subtype 3 |
|---|---|---|---|---|---|---|---|
| State Anxiety | 0.001 | 0.025 | 0.016 | 0.079 | 0.992 | 1 | 0.986 |
| Trait Anxiety | <0.001 | <0.001 | <0.001 | <0.001 | 0.971 | 0.975 | 1 |

**Supplementary Table 3:** *Post hoc* analyses of *n*-back activation. Group statistics were corrected for multiple comparisons by controlling the False Discovery Rate ($Q$<0.05). Pairwise contrasts are reported as *p*-values, and were adjusted via the Tukey method.

| | Pr(>F) | TD vs. Subtype 1 | TD vs. Subtype 2 | TD vs. Subtype 3 | Subtype 1 vs. Subtype 2 | Subtype 1 vs. Subtype 3 | Subtype 2 vs. Subtype 3 |
|---|---|---|---|---|---|---|---|
| Right Crus II | 0.043 | 0.490 | 0.195 | 0.215 | 0.031 | 0.036 | 1.000 |
| Left Anterior DLPFC | 0.043 | 0.633 | 0.070 | 0.213 | 0.017 | 0.056 | 0.989 |
| Dorsal Anterior Cingulate | 0.050 | 0.892 | 0.044 | 0.359 | 0.031 | 0.219 | 0.895 |
| Left Dorsal Frontal | 0.043 | 0.346 | 0.225 | 0.281 | 0.022 | 0.031 | 1.000 |
| Left Precuneus | 0.043 | 0.998 | 0.009 | 0.472 | 0.028 | 0.523 | 0.581 |
| Right Precuneus | 0.043 | 0.978 | 0.032 | 0.051 | 0.045 | 0.062 | 1.000 |

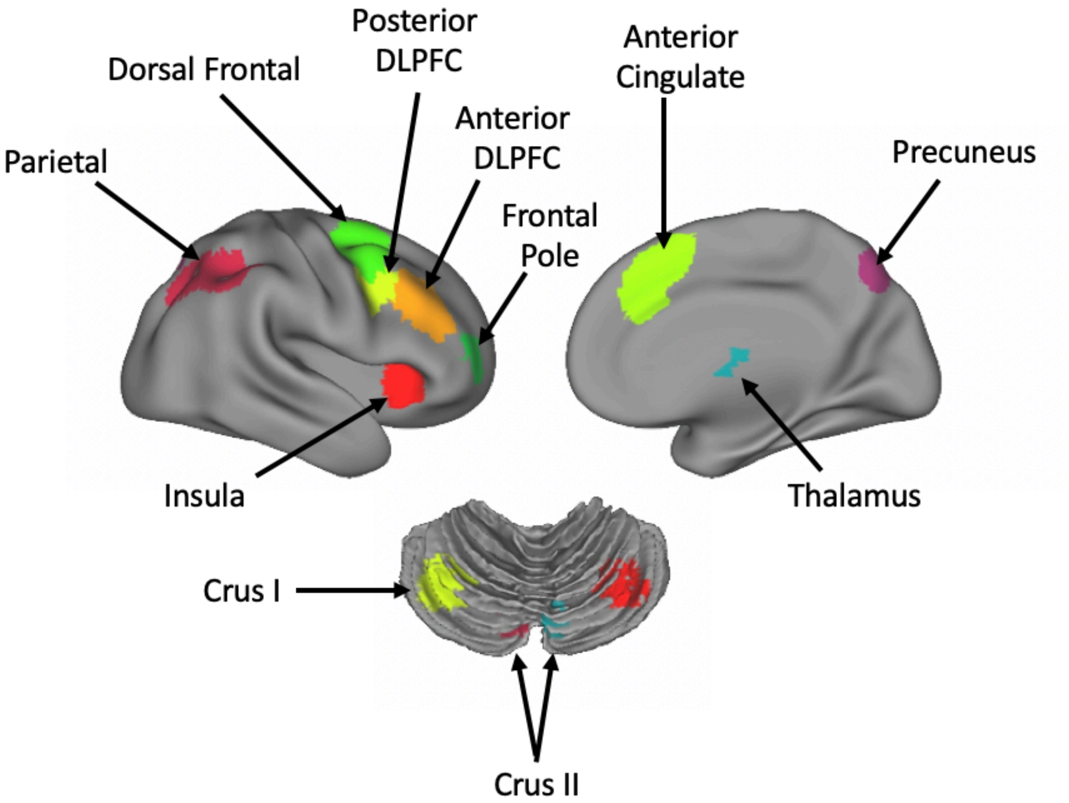


**Supplementary Figure 1:** Twenty-one functionally defined regions of interest. As described in previous work, the executive network was parsed into 21 functional regions of interest by applying a watershed algorithm to the map of the 2-back > 0-back contrast using an initial threshold of $z > 20$ (4).
